# A Novel ARNT-Dependent HIF-2α Signaling as a Protective Mechanism for Cardiac Microvascular Barrier Integrity and Heart Function Post-Myocardial Infarction

**DOI:** 10.1101/2023.03.12.532316

**Authors:** Karim Ullah, Lizhuo Ai, Yan Li, Lifeng Liu, Qin Zhang, Kaichao Pan, Zainab Humayun, Lin Piao, Albert Sitikov, Qiaozhu Su, Qiong Zhao, Willard Sharp, Yun Fang, David Wu, James K. Liao, Rongxue Wu

## Abstract

Myocardial infarction (MI) significantly compromises the integrity of the cardiac microvascular endothelial barrier, leading to enhanced leakage and inflammation that contribute to the progression of heart failure. While HIF2α is highly expressed in cardiac endothelial cells (ECs) under hypoxic conditions, its role in regulating microvascular endothelial barrier function during MI is not well understood. In this study, we utilized mice with a cardiac-specific deletion of HIF2α, generated through an inducible Cre (Cdh5Cre-ERT2) recombinase system. These mice exhibited no apparent phenotype under normal conditions. However, following left anterior descending (LAD) artery ligation-induced MI, they showed increased mortality associated with enhanced cardiac vascular leakage, inflammation, worsened cardiac function, and exacerbated heart remodeling. These outcomes suggest a protective role for endothelial HIF2α in response to cardiac ischemia. Parallel investigations in human cardiac microvascular endothelial cells (CMVECs) revealed that loss of ecHif2α led to diminished endothelial barrier function, characterized by reduced tight-junction protein levels and increased cell death, along with elevated expression of IL6 and other inflammatory markers. These effects were substantially reversed by overexpressing ARNT, a critical dimerization partner for HIF2α during hypoxia. Additionally, ARNT deletion also led to increased CMVEC permeability. Interestingly, ARNT, rather than HIF2α itself, directly binds to the IL6 promoter to suppress IL6 expression. Our findings demonstrate the critical role of endothelial HIF2α in response to MI and identify the HIF2α/ARNT axis as a transcriptional repressor, offering novel insights for developing therapeutic strategies against heart failure following MI.

## Introduction

Despite many therapeutic advancements, acute myocardial infarction (MI) remains the leading cause of death in Western countries (Khan, Hashim et al. 2020). The ischemic damage initiated by an intracoronary thrombus precipitates severe hypoxia, inflammation, and edema (Eltzschig and Eckle 2011). While many organs can tolerate mild to moderate edema, even small increases in interstitial fluid volume can significantly compromise cardiac function (Dongaonkar, Stewart et al. 2010). A fundamental cause of edema is an increase in vascular permeability. Current evidence supports the potential of vascular leakage inhibitors to improve cardiac outcomes by reducing hyperpermeability in cardiac microvascular endothelial cells (CMVECs) and limiting neutrophil infiltration. (Zhang, Kim et al. 2022) However, the mechanisms governing cardiac microvascular permeability remain underexplored as therapeutic targets in ischemic heart diseases.

The cellular response to hypoxia leads to the stabilization of Hypoxia-inducible Factors (HIFs), including Hif1α and Hif2α (HIF1A and HIF2A in humans), (Eckle, Kohler et al. 2008, Bautista, Castro et al. 2009, Du, Yang et al. 2020), which form heterodimeric complexes with aryl hydrocarbon receptor nuclear translocator (ARNT) to activate hypoxia-inducible genes (Maltepe, Schmidt et al. 1997, Koeppen, Lee et al. 2018, Nguyen, Zheng et al. 2021, Ullah and Wu 2021). Hif1α and Hif2α have been linked to several physiological and pathological processes that have a role in heart disease; However, the two isoforms appear to function independently and, in a cell-type–specific manner. Hif1α is ubiquitously expressed in all cell types and is known to have a cardioprotective role during ischemic preconditioning (Tekin, Dursun et al. 2010),(Sarkar, Cai et al. 2012, Zheng, Chen et al. 2021), whereas *Hif2*α*’s* expression is predominantly in endothelial cells (ECs) and its role in cardiac pathology is more complex and variable(Wiesener, Jurgensen et al. 2003). For example, although the cardiomyocyte-specific deletion of *Hif2*α (but not *HIF1*α) increased infarct sizes in a mouse model of ischemia-reperfusion injury (Koeppen, Lee et al. 2018), the inhibition of *Hif2*α reversed right heart failure and pulmonary arterial hypertension in rodents(Dai, Zhu et al. 2018). Pharmacological inhibition of HIF2a, such as PT2567, has been reported to improve pulmonary, vascular remodeling and hemodynamics in several preclinical models of established PH (Hu, Poth et al. 2019, Macias, Moore et al. 2021) Contrarily, the coronary microvasculature’s unique dilation in hypoxic conditions, as opposed to pulmonary artery constriction, raises questions(Sylvester, Shimoda et al. 2012, Marano, Wei et al. 2023, Zhao, Xiong et al. 2023), and this response is pivotal for enhancing oxygen supply to the myocardium. Consequently, the net effect of HIF2a inhibition on ischemic heart conditions, whether beneficial or detrimental, remains to be elucidated.

Although HIF1α is generally considered to be the primary mediator of the hypoxia response pathway(Eckle, Kohler et al. 2008, Bautista, Castro et al. 2009, Du, Yang et al. 2020), *Hif2*α expression has a protective role in kidney ischemia and myocardial ischemia-reperfusion injury (Kapitsinou, Sano et al. 2014), (Koeppen, Lee et al. 2018). Homozygous deletions of *Hif2*α are lethal during embryonic development(Licht, Muller-Holtkamp et al. 2006, Dunwoodie 2009, Skuli, Liu et al. 2009); However, mice with EC-specific deficiencies in *Hif2*α survive with varies aberrant phenotypes. (Skuli, Liu et al. 2009), (Gong, Rehman et al. 2015) Despite the evident role of Hif2α in myocardial disease, its specific functions in cardiac microvascular barrier function post-MI remain unclear. To address this knowledge gap, in this paper, we generated a mouse model with an inducible, EC-specific Hif2α knockout (ecHif2α–/–) to explore its role in this context. Our findings reveal that ecHif2α deficiency correlates with increased cardiac microvascular permeability and disruptions in endothelial barrier functionality, coupled with elevated IL6 expression in cultured ECs. Interestingly, the changes observed in our in-vitro studies were largely reversed by the overexpression of ARNT, and an ARNT point mutation that mimics its phosphorylation activity failed to reverse the effects. Furthermore, we found that HIF2α does not directly bind to the IL6 promoter. Instead, our results indicate that HIF2α negatively regulates IL6 expression through its binding partner ARNT, which directly binds to the IL6 promoter and regulates its expression. These findings suggest a novel mechanism by which Hif2α/ARNT regulates cardiac vascular function and inflammation in response to ischemic cardiac injury.

## Results

### Endothelial *Hif2***α** deletion impairs endothelial barrier function after MI

Our breeding program began with *Hif2*α*^flox/flox^* mice (C57BL/6 background; The Jackson Laboratory) and transgenic mice expressing tamoxifen-inducible Cre recombinase (CreERT2) under the control of the EC-specific VE-Cadherin promoter (Fig. 1A)(Alva, Zovein et al. 2006). After two generations, the *Hif2*α^flox/flox+*Cre*^ offspring were fed tamoxifen chow for two weeks to generate the experimental *ecHif2*α mice, and control assessments were conducted with *Hif2*α ^flox/flox^ mice fed tamoxifen chow and with *Hif2α*^flox/flox*+Cre*^ mice fed normal (tamoxifen-free) chow. (Supplemental Fig.1A-B). Mouse genotypes were verified via PCR (Fig. 1B), and the efficiency of ec*Hif2*α knockout induction was confirmed via analysis of Hif2α protein expression and *Hif2*α mRNA abundance in primary ECs isolated from *Hif2*α mice fed tamoxifen chow and *ecHif2*α^−/−^ mice upon completion of the feeding protocol (Fig.1C, D). The mice were phenotypically normal, with no significant changes in body weight, heart weight, blood pressure, or cardiac function (Supplemental Fig. 2-4). As neither control group showed any noticeable difference at baseline, for our study, we selected *Hif2*α^flox/flox^ mice as the control group, minimizing animal use without losing experimental rigor.

**Figure 1:**
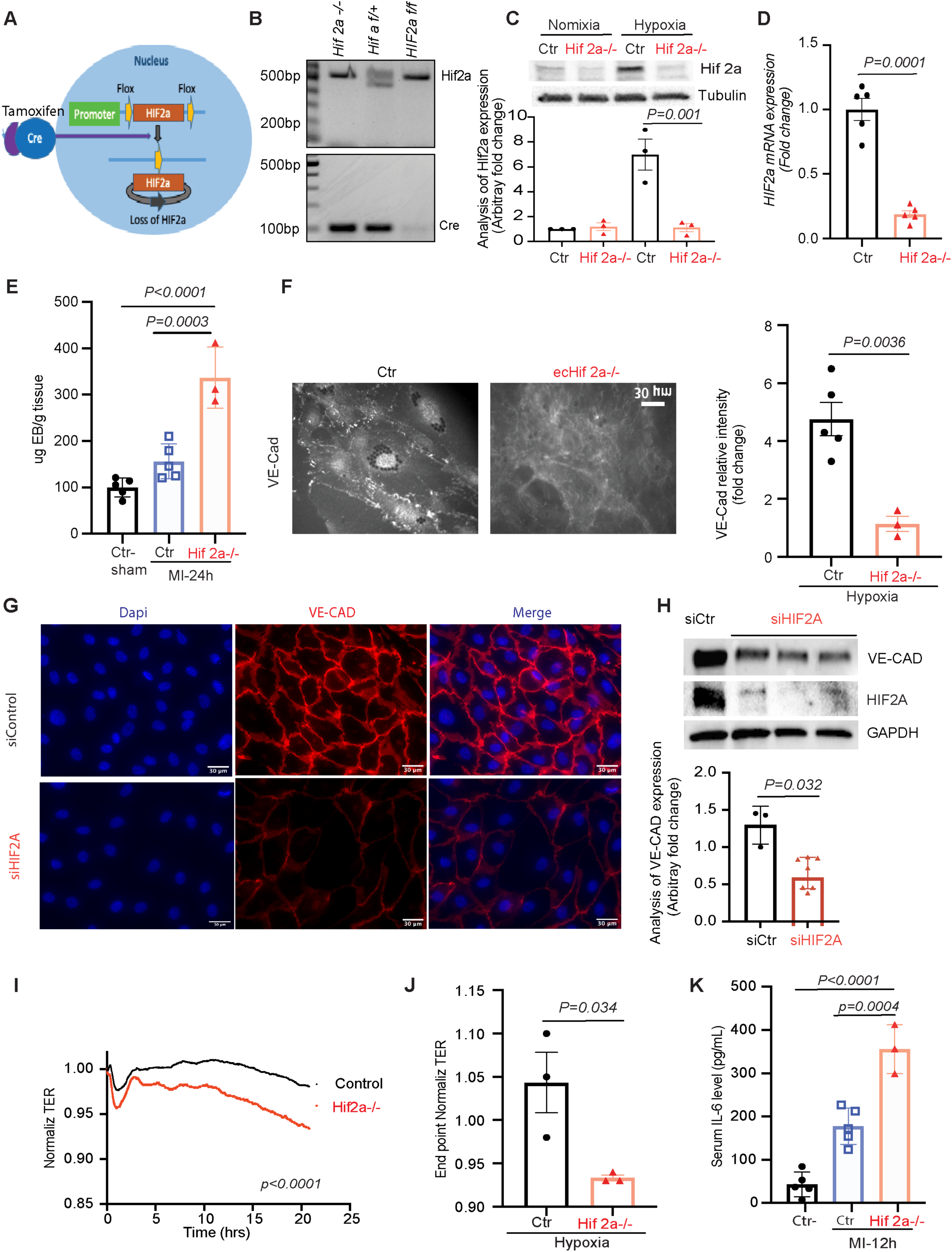
Endothelial *Hif2*α Deletion Impairs Barrier Function in MI. (A) Schematic breeding strategy to generate endothelial cells specific inducible *Hif2*α knockout mice. *Hif2*^flox/flox^ α were crossed with Cre-recombinase expressing mice under the control of VE-cadherin promoter. *Hif2*^flox/flox+CRE^ α and *Hif2^+/+ +CRE^*, *Hif2* flox/flox mice were grouped and fed with or without tamoxifen containing food for two weeks to induce CRE recombinase activity. After two weeks, the mice underwent experimental protocol. (B) Genotyping was determined by PCR. (C-D) WB analysis of cell lysates isolated from *Hif2*α^flox/flox+CRE^ and *Hif2*α mice (C). *Hif2*α mRNA expression analysis by RT-PCR confirmed the deletion of *Hif2*α gene (D). (E) Endothelial specific *Hif2*α deletion increases the permeability of heart. Quantification of Evans blue dye extravasation after 24h of MI. Control vs *Hif2*α^−/−^ MI mice p= 0.003, n=3. (F) Immunofluorescence staining of VE-Cadherin in control and *ecHif2*α*^−/−^* EC. (Left panel) Quantification of VE-Cadherin intensity in control and *ecHif2*α*^−/−^* cells, control vs *ecHif2α^−/−^* α p=0.0036, n=3-5 (G) Immunofluorescence staining of VE-Cadherin in primary Human Microvascular Endothelial Cells (HMVEC) transfected with or without si*Hif2*α and treated with 1% oxygen for overnight. The nuclei were stained DAPI (Blue), Ve-Cadherin (Red), Scale bar: 30 µm. (H) WB analysis of whole cell lysates from HMVEC transfected with or without scrambled or si*Hif2*α and treated with 1% oxygen for 16h. Quantification of VE-Cadherin protein levels (n=4) in lower panel. (I) Analysis of ECIS assays for *ecHif2*α*^−/−^* and wt EC cells. 90-100 % confluent monolayers of wt and *ecHif2*α*^−/−^* cells were cultured in collagen coated ECIS 8W10E+ arrays and treated with 1mM of DMOG to induce hypoxia. TER of cells was measured in Ohms using electric cell-substrate impedance sensing (ECIS) technology. Measurements were taken continuously at 3-minute intervals. (J) Analysis of endpoint normalized TER for ECIS assays. Results are shown as mean ± SEM for n=4 independent experiments. (K) ELISA measurement of plasma IL6 levels in MI induced *ecHif2*α*^−/−^* and control mice, n=3. Results are shown as mean ± SEM.

MI was induced in *ecHif2*α*^−/−^* mice and in Control mice via ligation of the left anterior descending (LAD) coronary artery, and another group of *ecHif2*α*^−/−^* mice (the Sham group) underwent all surgical procedures for MI induction except coronary artery ligation (Supplemental Fig. 5). Evans Blue dye was injected into the tail veins of mice 12 hours after MI induction or Sham surgery, and hearts were harvested 30 minutes later after Evans blue injection. Measurements of Evans blue extravasation were significantly greater in *ecHif2*α^−/−^ hearts than in the hearts of Control or Sham mice (Fig. 1E), but significant differences were not observed between the two groups for organs under normoxic condition (Supplemental Fig. 6). Furthermore when cells were cultured under hypoxic conditions, VE-cadherin was significantly less abundant in CMVEC isolated from *ecHif2*α^−/−^ hearts than in Control ECs (Fig. 1F) and in human microvascular endothelial cells (hMVECs) (Fig. 1G-H) as well as HUVEC (Supplemental Fig. 7) transfected with *Hif2*α siRNA than in control siRNA transfected cells. In addition, the expression of Zonula occludens-1 (ZO-1), a tight junction protein that plays a critical role in endothelial barrier function, was also reduced in hMVECs with *Hif2*α deficiency under hypoxia condition. (Supplemental Fig. 8). In ex vivo assays using the ECIS system, mCMVECs from ecHif2α–/– mice had lower transendothelial resistance (TER) compared to controls, the difference that was pronounced under DMOG-induced hypoxic conditions (Fig. 1I-J) but absent under normoxia (Supplemental Fig. 9).

In ex vivo measurements using the ECIS system, transendothelial resistance (TER) was significantly lower in monolayers of mouse cardiac microvascular endothelial cells (mCMVECs) from ecHif2α–/– mice than in control mCMVEC monolayers. Since the ECIS system cannot serve as a hypoxia chamber, we used DMOG (dimethyloxalylglycine) to mimic a hypoxic condition. Under these DMOG-induced hypoxic conditions, the differences were evident (Fig. 1I-J). However, no differences were detected under normoxic conditions (Supplemental Fig. 9). These results indicate that endothelial-specific loss of *Hif2*α expression increased vascular permeability in infarcted hearts, reduced junctional protein expression in cultured ECs, and impaired barrier function in mCMVEC monolayers. Notably, 12 hours post-MI, compared with controls, ecHif2α–/– mice exhibited elevated serum IL6 levels, implicating Hif2α in modulating inflammatory responses to cardiac injury, specifically through IL6 (Fig. 1K). In contrast, no significant changes in TNFα or IL1b levels were noted (Supplemental Fig. 10).

### Loss of ec *Hif2***α** increased mouse mortality and cardiac remodeling following myocardial infarction

Cardiac function was assessed using echocardiography (Fig. 2A) at baseline (prior to MI induction), as well as on days seven and 28 post-MI. Initial evaluations of left ventricular ejection fraction (EF, Fig. 2B), fractional shortening (FS, Fig. 2C), and cardiac output (Fig. 2D) revealed no significant differences between ecHif2α–/– and Control mice. However, by Day 7, EF had notably decreased in ecHif2α–/– mice, and by Day 28, reductions were evident across all three parameters in this group. Examinations of hearts harvested on Day 28 demonstrated that the absence of ecHif2α led to pronounced heart enlargement (Fig. 2E), increased myocardial fibrosis, and heightened inflammatory cell infiltration (Fig. 2F). The expression of Col13a1 and Lox3, genes involved in fibrotic processes, was markedly increased (Fig. 2G and Supplemental Fig. 11A), aligning with enhanced pathway activity related to fibrosis and heart failure, as depicted in Figure 3. Corresponding to these alterations, heart failure markers, such as BNP and NPR3, were significantly higher in ecHif2α–/– mice, suggesting a molecular underpinning for the observed phenotypic changes (Figs. 2H-I). Critically, survival rates substantially declined in ecHif2α–/– mice compared to their Control (Fig. 2J), demonstrating the potential of ecHif2α as a key factor in survival post-MI.

**Figure 2:**
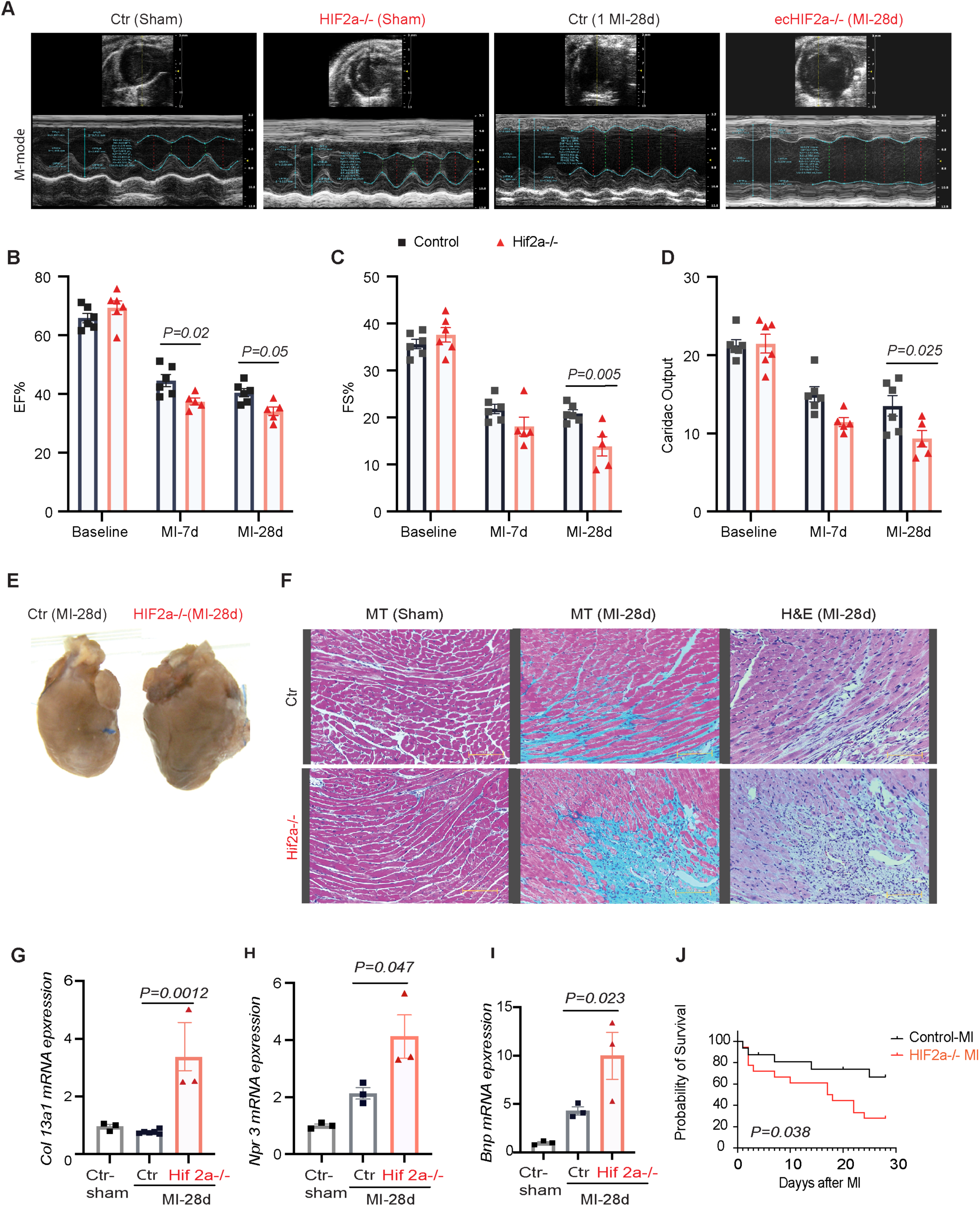
Impact of Hif2α Deletion on Cardiac Function and Fibrosis Post-MI. (A) Comparison of cardiac function in Control and ecHif2α–/– mice, post-MI at 7 and 28 days. Representative M-mode echocardiography images are presented. (B-C) Evaluation of Ejection fraction (EF) and Fractional shortening (FS) at various time points post-MI. n=6 (D) Cardiac output measurement. (E) Representative images of heart 28 days post-MI. (F) Histological analysis of heart sections stained with hematoxylin-eosin and Masson trichrome. (G-I) qPCR assessment of mRNA levels of select genes *Col13a1, Npr3* and *Bnp* in heart tissues. Data represented as meanL±LSEM, n=3. (J) Kaplan-Meier survival curve post-MI. n=34, p=0.05.

**Figure 3:**
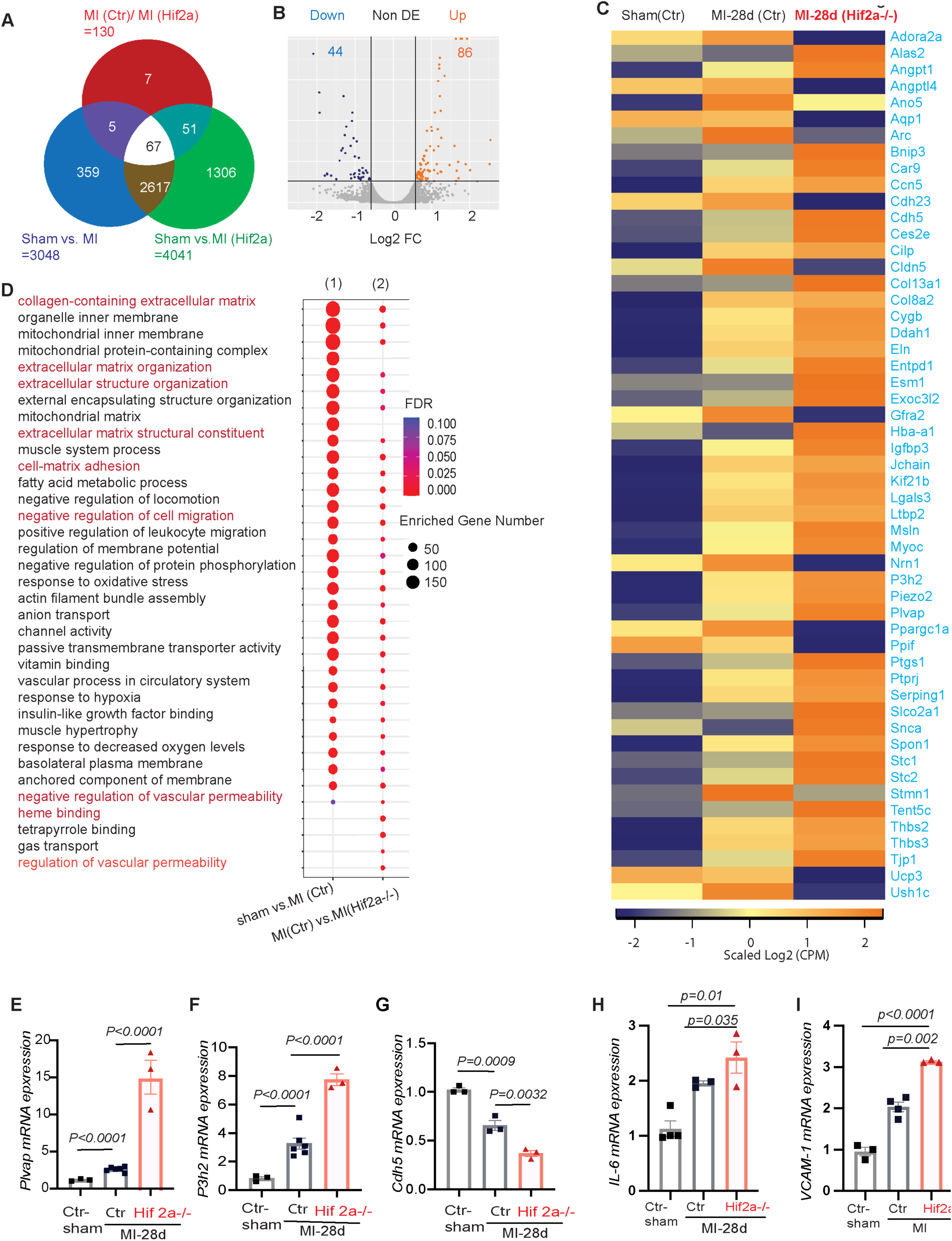
RNA-Seq Insights into Gene Expression Changes in Heart Post-MI. (A) Venn diagram showing the overlap of MI induced genes between controls and *ecHif2*α heart tissues. (B) Volcano map of differentially expressed genes between controls and *ecHif2α^−/−^* α heart tissues. Orange dots represent the upregulated genes and dark blue dots represent the downregulated genes. (C) Heatmap of differentially expressed genes in MI induced controls and *ecHif2*α*^−/−^* heart tissues. (D) KEEG pathway enrichment analysis of differentially expressed genes in controls and *ecHif2*α*^−/−^* samples shows dysregulation of multiple signaling pathways associated endothelial barrier function. (E-I) mRNA expression analysis of *Plvap* (E), *P3h2* (F), *Cdh5* (G), *Il-6*(H) and *Vcam-1* (I) in controls and *ecHif2*α*^−/−^* samples after 28 days of MI. Data are presented as meanL±Ls.d., n=3.

### Differential gene expression profiles in *ecHif2***α***–/–* mice reveal altered vascular dynamics post-myocardial infarction

Heart tissues subjected to bulk RNA sequencing 28 days post-myocardial infarction (MI) or sham operation unveiled distinct changes in gene expression in ecHif2α–/– mice compared to controls. Of 130 differentially expressed genes (DEGs), 86 were upregulated, in which many implicated in increasing vascular permeability and promoting fibrosis. (Fig. 3A, Supplemental Tables 1-4). Conversely, the remaining 44 downregulated DEGs included many of those that promote vascular barrier function (Fig. 3B-C). These observations were supported by pathway enrichment analysis, which indicated that genes involved in vascular permeability, cell migration, and cell-matrix adhesion were significantly enriched in the DEGs (Fig. 3D). Moreover, measurements of mRNA abundance confirmed that regulators of vessel-permeability and collagen synthesis (*plvap* and *P3h2* respectively) were more highly expressed. In contrast, key regulators of endothelial barrier function (such as cdh-5, tjp-1, and angpt-2) exhibited diminished expression compared to Control hearts (Fig. 3E-G and Supplemental Fig. 11 B, C). Notably, the *ecHif2*α deletion was also associated with a significant increase in mRNA levels for the inflammatory IL6 and VCAM-1 (Fig 3H, I), while ICAM and e-selectin expression tended to be greater (though not significantly) in infarcted *ecHif2*α hearts than infarcted Control hearts (Supplemental Fig. 12) which is consistent with our observation that the MI-induced cardiac infiltration of inflammatory cells was more remarkable in *ecHif2*α mice than in control mice.

### *Hif2*α deficiency impairs angiogenetic activity and promotes apoptosis in cultured Endothelial cells (ECs) under hypoxia

Because vascular permeability and angiogenesis are closely linked (Khurana, Simons et al. 2005, Fahey and Doyle 2019), we investigated whether the aberrations in vascular barrier function associated with *ecHif2*α deletion were accompanied by changes in the angiogenic activity of ECs. Using a tube formation assay on Hif2α-silenced HUVECs, we observed a marked decrease in the complexity of the angiogenic network under hypoxic conditions, as evidenced by fewer nodes and junctions compared to control cells (Fig. 4A). This impairment was quantifiable; both the meshwork and the total length of the tubes formed were considerably reduced in Hif2α-deficient cells in both normal and low-oxygen environments (Figs. 4B-E).

Further analysis under normal oxygen conditions showed no significant differences in cellular health or apoptosis between Hif2α-deficient HUVECs and controls (Figs. 4F-H). However, under low oxygen, a condition mimicking the post-MI environment, Hif2α-silenced HUVECs exhibited a marked increase in early-stage These observations collectively imply that Hif2α is essential for maintaining angiogenic processes and endothelial cell survival, particularly in hypoxic conditions like those found in infarcted myocardial tissue.

### Restoration of endothelial barrier integrity by ARNT overexpression in Hif2**α**-Deficient endothelial cells

Given the heterodimeric function of Hif2α with ARNT(Maltepe, Schmidt et al. 1997), we investigated whether ARNT overexpression could counteract the deleterious effects of Hif2α deficiency. Experiments were conducted in HMVECs and HUVECs that were transfected with siHif2α or control (scrambled) siRNA, and co-transfected with vectors coding for wild-type ARNT (ARNT^WT^) or an ARNT mutant)(Bourner, Muro et al. 2022). The nuclear localization of overexpressed ARNT was confirmed and behaved as anticipated. (Supplemental Fig. 13). Notably, overexpression of ARNT^WT^, but not ARNT^mut^, reduced hypoxia-induced apoptosis (Fig. 5A) and restored VE-cadherin expression to normal levels in si*Hif2*α-transfected HMVECs (Fig. 5B).

**Figure 4:**
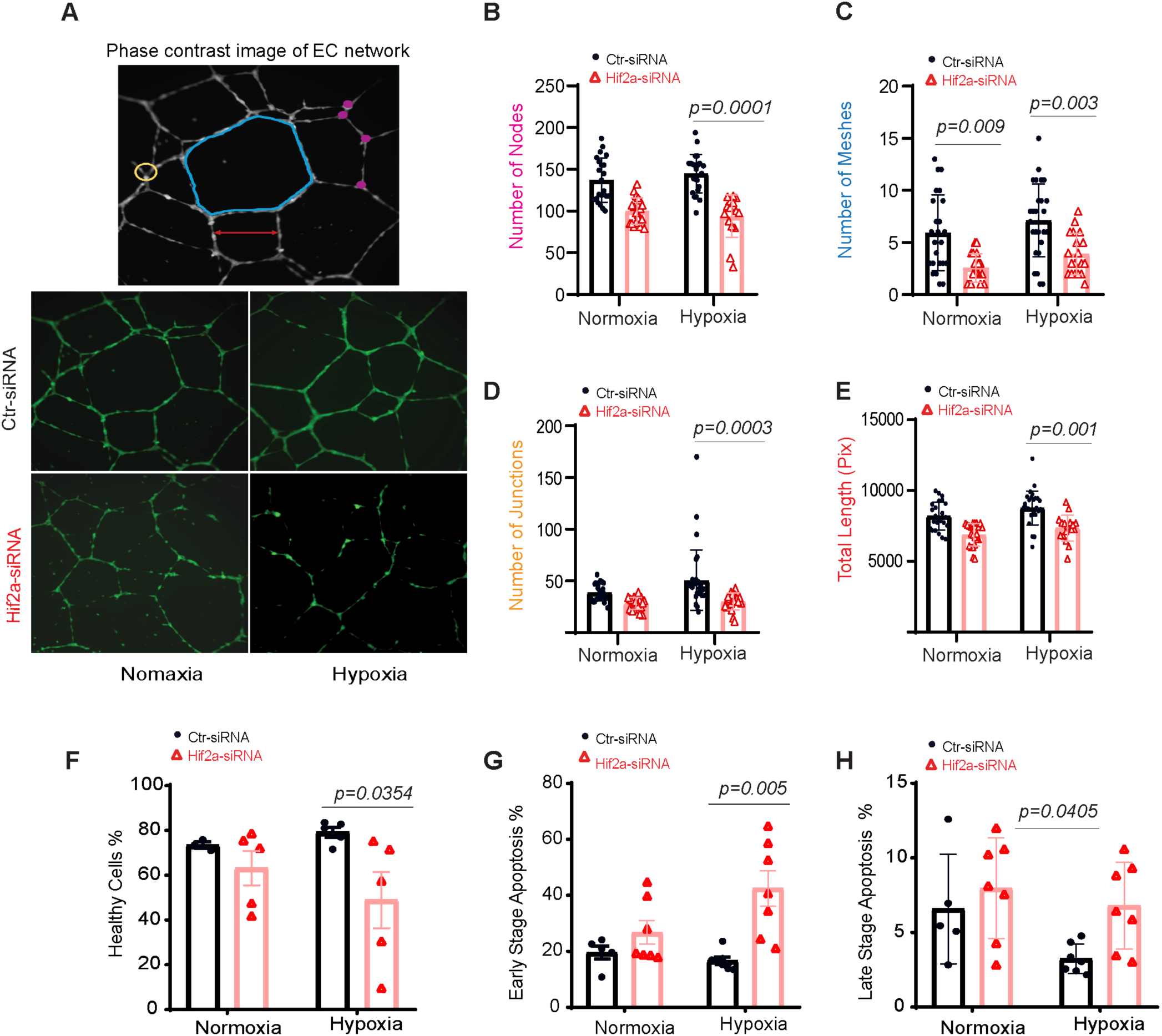
Hif2α Deletion Impacts Angiogenesis and Increases Endothelial Cell Apoptosis. (A) HUVEC network visualization via Image J. Top: Meshes (blue), nodes (pink), junctions (yellow), and segment lengths (red). Bottom: Capillary-like structures from 6 to 16h post-seeding. (B-E) Metrics for nodes (B), meshes (C), junctions (D), and total length (E). n=24. (F-H) Apoptosis in HUVECs post Hif2α silencing. Cells were transfected with Hif2α siRNA or control and subjected to 1% oxygen or normoxia. Flow cytometry assessed healthy (F), early apoptotic (G), and late-apoptotic cells (H) using 7-AAD labeling. Analysis via two-way ANOVA with Sidak’s test. n=4 or 5.

**Figure 5:**
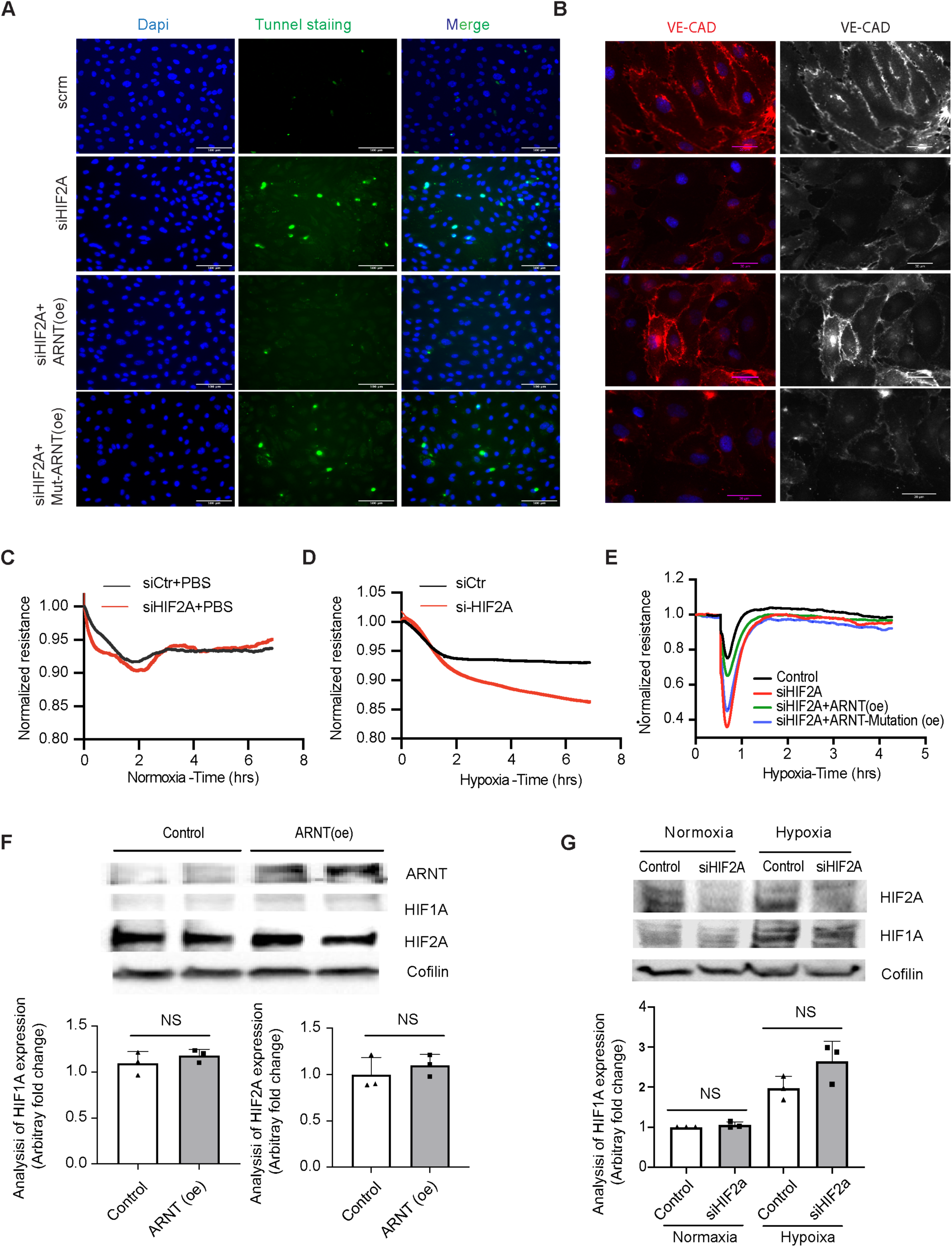
Interplay of Hif2α Deletion and ARNT Variants in Hypoxic HMVEC Barrier Dysfunction. (A) Tunnel staining in HMVECs transfected with siHif2α and overexpressed with either ARNT-WT or Mutant-ARNT post 1% O₂ exposure for 16h, n=8. (B) Immunofluorescence of VE-Cadherin in HMVECs with Hif2α silencing and either ARNT-WT or Mutant-ARNT overexpression, n=3. (C-D) ECIS post Hif2α silencing and treatment with PBS (C) or DMOG (D), n=4. (E) ECIS in HMVECs post Hif2α silencing and overexpression of either ARNT-WT or Mutant-ARNT, followed by 16h DMOG exposure, n=4. (F-G) Western blots indicating ARNT, HIF1A, and HIF2A levels post varied transfections. Coflin as control, n=3. Data represented as mean ± SEM; NS indicates no significance.

Using Electric Cell-substrate Impedance Sensing (ECIS), we assessed barrier function in HUVEC monolayers. Barrier integrity was compromised under hypoxic conditions induced by dimethyloxalylglycine (DMOG), a hypoxia mimetic, rather than at normoxia (Fig. 5C-D). (Note: The ECIS system cannot directly induce hypoxia; hence, DMOG is employed to generate cellular hypoxia to assess cell barrier integrity using the system, as mentioned before.) Measurements of minimum resistance, which occurred 40 minutes after hyperpermeability was induced via treatment with thrombin, were much greater in HMVEC monolayers when the cells were transfected with both si*Hif2*α and ARNT^WT^ than with si*Hif2*α alone or si*Hif2*α and ARNT^mut^ (Fig. 5E). Subsequent evaluation of HIF1α protein levels under hypoxia post-ARNT overexpression or Hif2α knockdown revealed that Hif1α concentrations were largely stable, suggesting that the observed phenotypic changes were not due to compensatory activity by the Hif1α isoform (Fig. 5F-G).

Furthermore, treatment with the HIF2α inhibitor, PT2567, led to a paradoxical increase in IL6 and IL1b mRNA levels, enhancing IL6 expression detrimentally affecting VE-cadherin (VE-CAD) expression, a crucial component of endothelial barrier integrity. Remarkably, these effects were reversed by IL6 knockdown via siRNA (Fig. 6A-B), reinforcing the link between HIF2α activity and endothelial barrier function (Fig. 6C).

**Figure 6:**
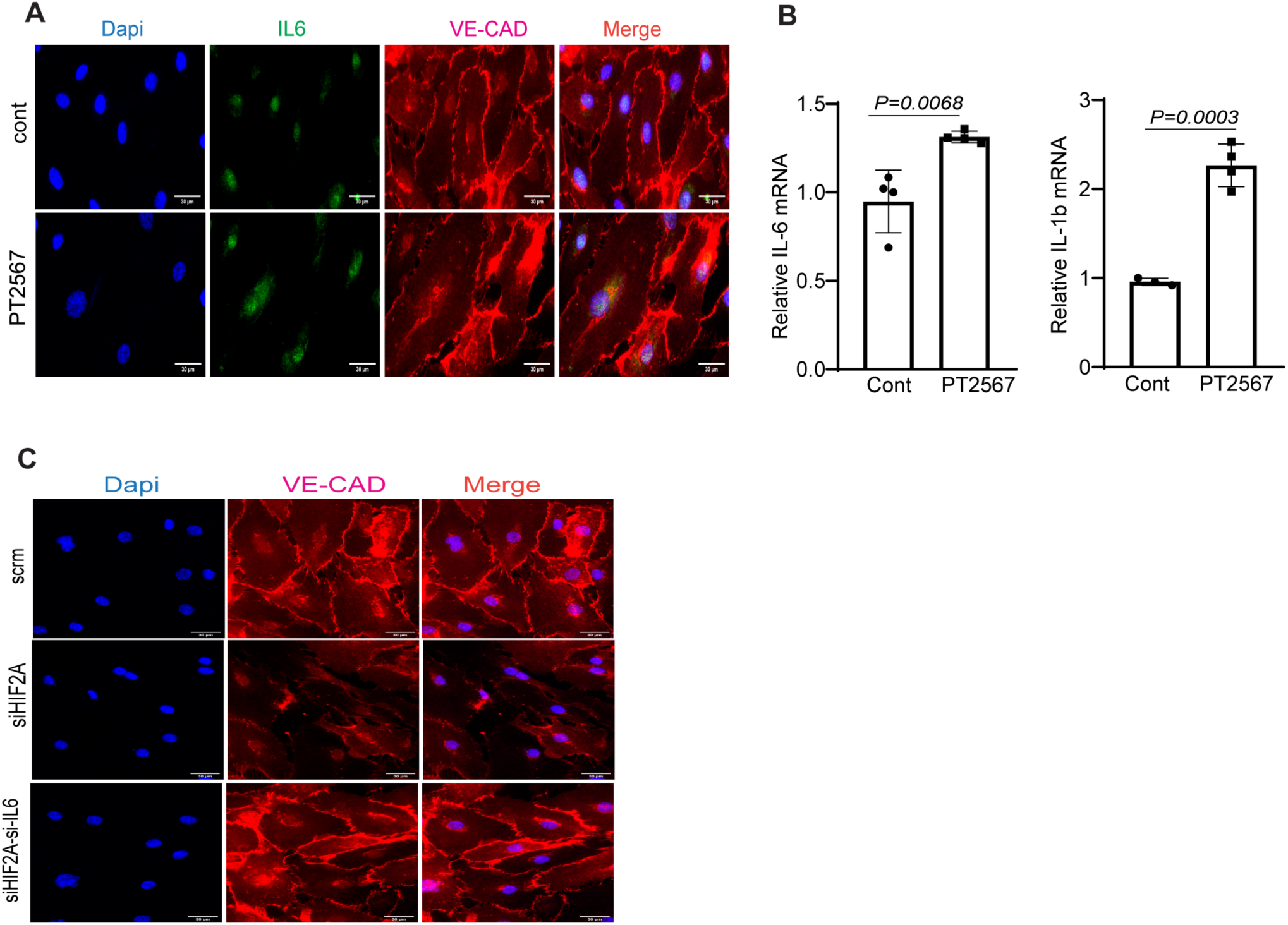
(A) HMVECs were treated with 10uM of PT2567 and exposed to 1% hypoxia for 16h. IL-6 and IL-1b mRNA expression were analyzed by RT-PCR. (B) Immunofluorescence staining of IL-6 and VE-Cadherin in PT2567 treated HMVEC. The respective stains were Nucleus Dapi, IL-6 (Green), and VE-Cadherin (Red). The scale bar denotes 30μM.(C)

### HIF2**α** regulates IL6 expression through direct binding of ARNT to the IL6 promoter

Delving into the potential mechanisms, we focused on how ARNT might counteract the inflammation increase associated with Hif2α reduction. We found that the absence of ARNT alone increased the levels of inflammatory markers in HMVECs, including IL6 protein expression and ICAM, VCAM-1 mRNA levels (Fig. 7 A-D). We also observed that ARNT overexpression reverses siHif2α-Induced IL6 Expression in HMVECs. (Fig. 8A). Similar results were observed in human aortic endothelial cells (HAECs) subjected to hypoxic conditions, where co-transfection with siHif2α and ARNT^WT^ plasmid led to lower IL6 mRNA and protein levels compared to siHif2α alone (Fig. 8B-F).

**Figure 7:**
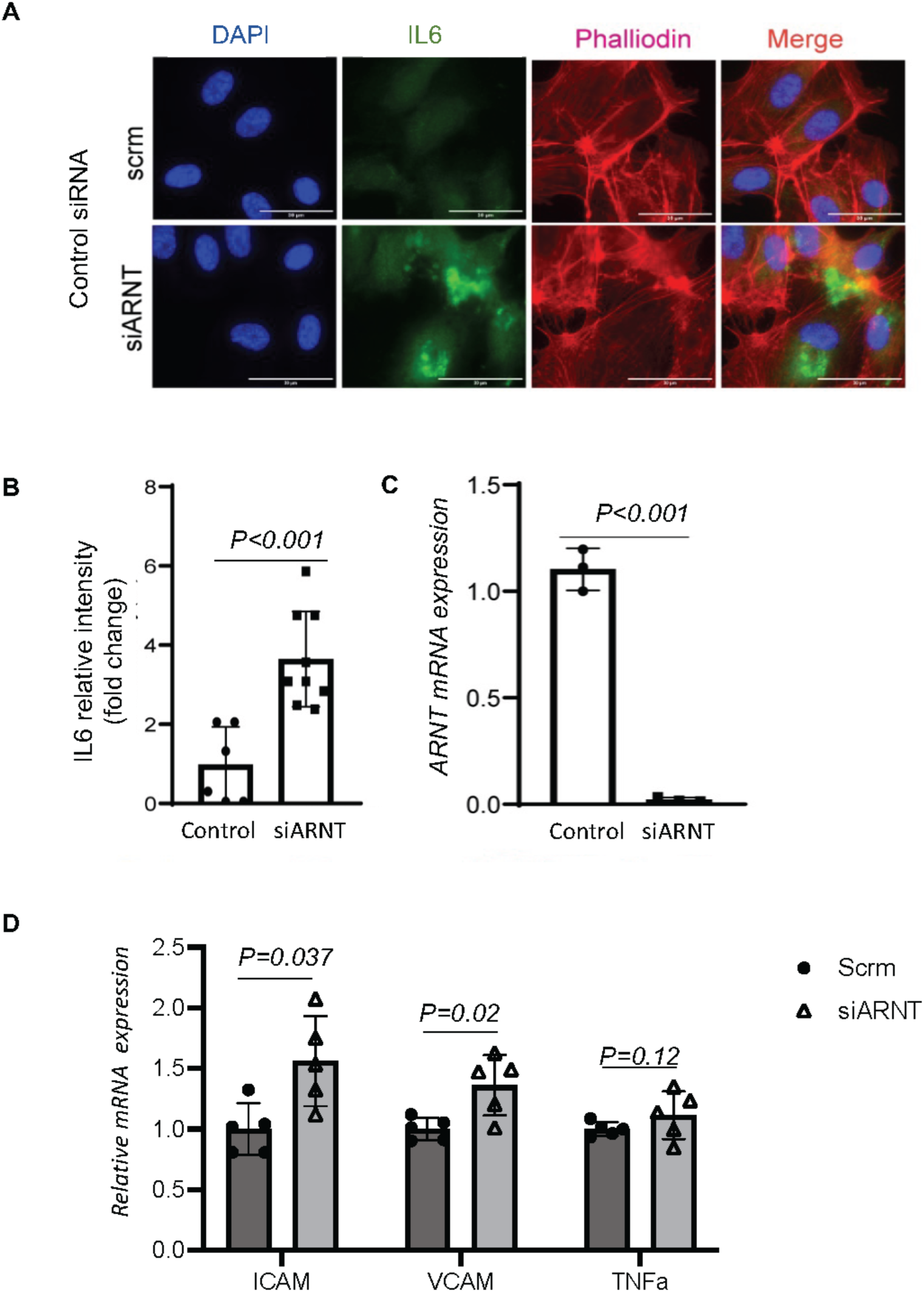
**C** In HMVECs, Ve-Cad expression is revealed through immunofluorescence staining. Cells underwent siRNA-mediated knockdown of IL6 and HIF2α for a span of 48-72 hours and were subsequently exposed to 1% hypoxia throughout the night. Staining colors are as follows: Ve-Cad in red and DAPI in blue. Supplementa Fig 7: Analysis of IL6 Expression and Intensity Following ARNT Deletion in HMVECs. HMVECs underwent siRNA-mediated ARNT knockdown for a period of 48-72 hours, followed by an exposure to 1% hypoxia overnight. A) Immunofluorescence staining illustrates IL6 expression highlighted in green, with the nuclei counterstained in blue using DAPI. The scale bar denotes 30μM. B) Quantitative analysis of IL6 intensity from the immunofluorescence staining. C) Evaluation of ARNT mRNA levels post-siRNA treatment. Data are presented as mean ± SEM (n=3-4). Supplemental Fig14: Evaluation of Inflammatory Marker Expression Post-ARNT Deletion. Human Microvascular Endothelial Cells (HMVECs) underwent siRNA-mediated ARNT knockdown for 48-72 hours and were subsequently subjected to 1% hypoxia overnight. Control siRNA treatment is represented as scramble (Scrm). Assessed genes encompass the Intercellular Adhesion Molecule (ICAM), Vascular Cell Adhesion Molecule (VCAM), and Tumor Necrosis Factor Alpha (TNFα). Data are denoted as mean ± SEM (n=3-4).

**Figure 8:**
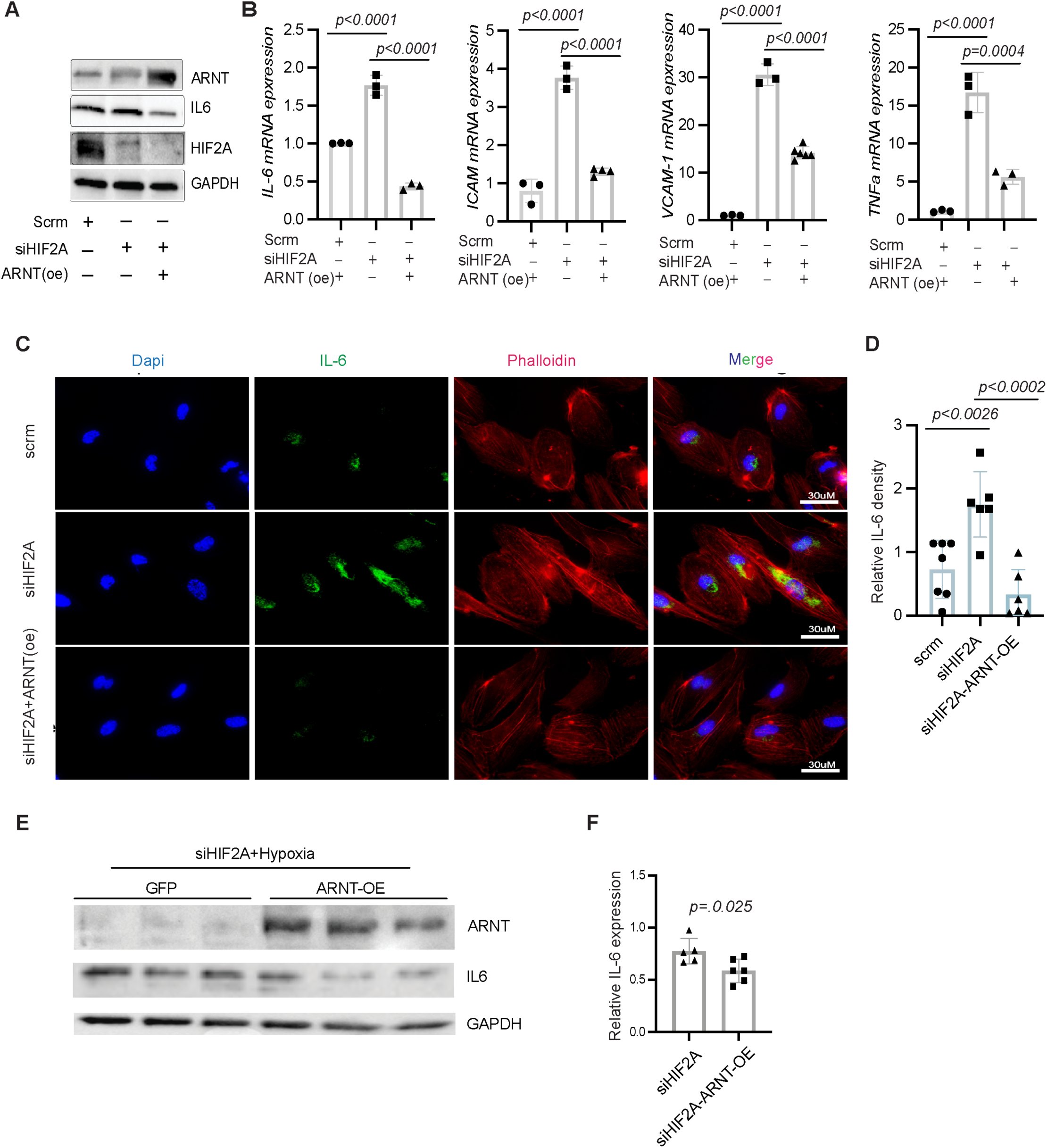
Impact of ARNT Overexpression on Hypoxic Inflammatory Response in Hif2α Silenced Human Cardiac Endothelial Cells. (A) Western blot analysis of ARNT, IL6, and Hif2α in HMVECs transfected with Hif2α siRNA and either overexpressed with ARNT or an empty vector under hypoxic conditions. (B) RT-PCR determination of mRNA expression levels of IL-6, ICAM, VCAM-1, and TNFα in HAECs. 18s rRNA serves as an internal control. (C) Immunofluorescence of IL-6 in HAECs; phalloidin staining highlights cellular skeletal structure. (D) Quantitative analysis of IL-6 fluorescence intensity in HAECs. (E) Western blot of ARNT and IL6 in HAECs transfected with Hif2α siRNA, either overexpressed with ARNT or GFP serves as a control and exposed to 1% O₂ for 16 hours. Data represented as mean ± SEM.

Experiments in HEK293 cells equipped with an IL6-luciferase promoter-reporter led to a significant drop in luciferase activity in hypoxic conditions (Fig. 9A-B). Conversely, transfection with siHif2α alone bolstered luciferase activity. However, when co-transfected with siHif2α and ARNT^WT^ overexpression, the luciferase activity remained the same to control cells and was notably diminished compared to the siHif2α-transfected cells or those co-transfected with siHIF2α and ARNT^mut^ (Fig. 9C). Chromatin immunoprecipitation followed by quantitative PCR (ChIP-qPCR) further demonstrated direct binding of ARNT, but not Hif2α, to the IL6 promoter (Figures 9D-E), indicating a potential role for ARNT in the regulation of IL6 in the context of HIF2a signaling.

**Figure 9:**
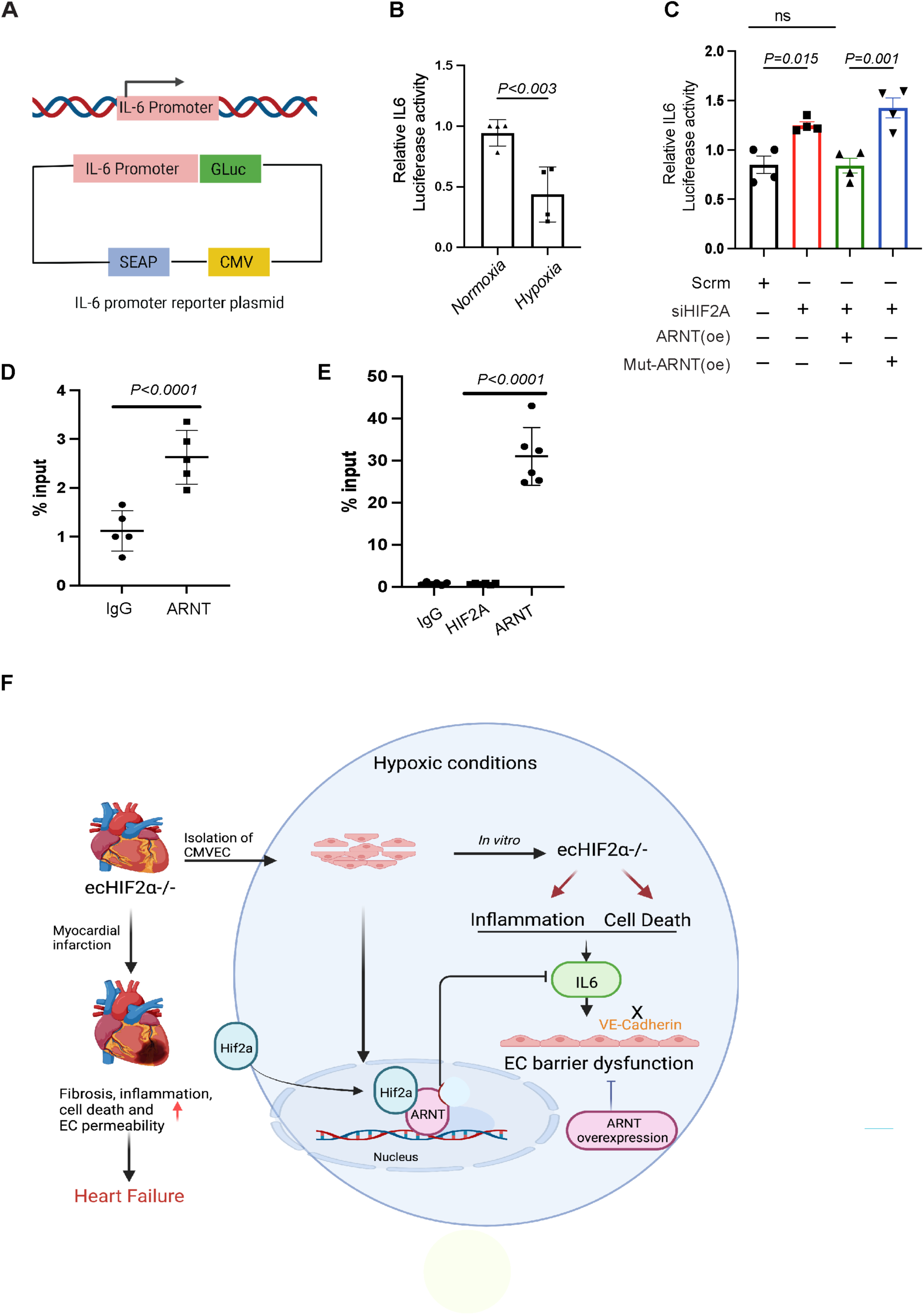
Direct Inhibition of IL6 Transcription by Hif2α/ARNT Heterodimers through DNA Binding. (A) Schematic representation of the IL-6 promoter reporter construct. (B) Analysis of normalized IL-6 promoter activities in HEK293 cells. Cells transfected with the human IL6 dual-reporter promoter construct were either exposed to ambient oxygen levels or subjected to 1% oxygen for 24 hours (n=4). (C) Protocol followed in HEK293 cells: Initial transfection was with or without controls and Hif2α siRNA. Post 24 hours, a dual-reporter vector containing the human IL6 promoter was introduced and allowed to incubate overnight. Concurrently, cells underwent transfection with lentiviral vectors encoding ARNT-WT or ARNT-Mut. This was followed by a 1% oxygen treatment for 24 hours prior to sample collection (n=4). (D-E) ChIP-qPCR analysis of IL6. HEK293 cells either underwent a 1% oxygen treatment or were transfected with an ARNT-encoding lentivirus before exposure to 1% oxygen. Chromatin immunoprecipitation used the specified antibodies. Data represented as mean ± SEM F) Diagram of Mechanism of Hif2α/ARNT-mediated regulation of cardiac endothelial barrier function and cardiac function during myocardial infarction.

## Discussion

We demonstrate a novel aspect of Hif2α signaling that is critical for maintaining endothelial barrier integrity, particularly in the condition of myocardial infarction (MI). Our study reveals that endothelial-specific loss of Hif2α significantly increases vascular permeability and exacerbates cardiac remodeling by impairing junctional protein expression, such as VE-cadherin and Zonula occludens-1. Additionally, the elevated serum IL6 levels and reduced angiogenic response in Hif2α-deficient endothelial cells emphasize its pivotal role in modulating inflammation and angiogenesis under hypoxic conditions. Importantly, the suppression of IL6 transcription through ARNT overexpression highlights the therapeutic potential of the HIF2α/ARNT axis in improving recovery after MI. Collectively, our observations are consistent with previous reports linking microvascular dysfunction to a variety of pathological processes, including inflammation, apoptosis(3, 33),(34-36), and impaired angiogenesis(37, 38), and suggest that Hif2α/ARNT heterodimers are crucial regulators of endothelial barrier integrity and function in ischemic hearts.

### Impact on Endothelial Barrier Function and Cardiac Health

The use of inducible endothelial cell-specific Hif2α knockout mice (ecHif2α–/–) allowed for a targeted investigation into the role of Hif2α within the endothelium in adult mice, revealing that its absence led to increased vascular permeability and reduced expression of junctional proteins like VE-cadherin and ZO-1. VE-cadherin, being a primary tight junction protein, is crucial in maintaining endothelial barrier integrity(Duong and Vestweber 2020). The perturbed integrity of the endothelial barrier in ecHif2α–/– mice was further validated by the pronounced Evans Blue extravasation in infarcted hearts and decreased transendothelial resistance (TER) observed with the ECIS system. These findings align with the current understanding that endothelial junctional disruption correlates with increased vascular permeability in pathological conditions (Aghajanian, Wittchen et al. 2008, Lee 2015). The decline in endothelial barrier integrity leads to cardiac vascular leakage and cardiac edema, likely facilitating the infiltration of inflammatory cells and substances into the myocardium, thereby worsening cardiac injury and remodeling. These changes were mirrored by a significant decline in cardiac function in ecHif2α–/– mice, particularly noted in the deterioration of ejection fraction, fraction shortening, and increased myocardial fibrosis. However, study also reported that increased microvascular permeability in heart Induced diastolic dysfunction independently of inflammation, fibrosis, or cardiomyocyte dysfunction (Abelanet, Camoin et al. 2022)

### Endothelial HIF2α deletion and cardiac fibrosis post-MI

Our study elucidates a notable link between endothelial HIF2α deletion and a cascade of events, including compromised cardiac endothelial barrier function, increased inflammation, and augmented cardiac fibrosis. However, the precise mechanisms by which endothelial HIF2α deletion contributes to cardiac fibrosis following MI are likely multifaceted. The leakage of pro-inflammatory factors due to endothelial dysfunction likely contributes directly to cardiomyocyte damage and subsequent fibroblast activation. Additionally, elevated IL-6 levels may promote the transition of fibroblasts into collagen-producing myofibroblasts, exacerbating cardiac stiffening and fibrosis. The potential role of HIF2α in regulating endothelial-to-mesenchymal transition (EndMT) suggests a complex mechanism, although direct evidence remains speculative. Future studies are essential to delineate the specific molecular pathways influenced by HIF2α in endothelial cells, particularly those regulating EndMT.

### Role of Hif2α in Angiogenesis and Inflammatory Modulation

Interestingly, our study extends the known functions of Hif2α to include its involvement in angiogenesis under hypoxic conditions. The observed reduction in angiogenic activity, as demonstrated by impaired tube formation in Hif2α-silenced HUVECs, suggests that Hif2α is essential for the angiogenic response following hypoxic stress. This is particularly critical in the post-MI healing process, where angiogenesis is crucial for tissue repair and recovery. Furthermore, Hif2α appears to regulate inflammatory processes differently in endothelial cells compared to cardiac myocytes, as demonstrated by the negative regulation of IL6.(Wu, Hu et al. 2021). The elevated IL6 levels in ecHif2α–/– mice post-MI could be directly linked to the observed increases in vascular permeability and inflammation. Numerous studies have shown that HIF activation is associated with inflammation (Meneses and Wielockx 2016, Watts and Walmsley 2019, Rajendran, Schonfeld et al. 2020), and that inflammation increases vascular permeability by destabilizing endothelial junctional proteins (He, Chen et al. 2014, Chistiakov, Orekhov et al. 2015, Castellon and Bogdanova 2016), (Gong, Rehman et al. 2015). IL6 is a prominent regulator of both inflammation (Yang, Fu et al. 2022) and vascular permeability(Alsaffar, Martino et al. 2018). Although the role of IL6 induction in the ECs of infarcted hearts has yet to be thoroughly characterized (van den Oever, Raterman et al. 2010, Su, An et al. 2018, Claesson-Welsh, Dejana et al. 2021), inhibition of IL6 attenuates inflammation and prevents endothelial barrier disruption in retinal ECs(Valle, Dworshak et al. 2019). Here, we provided direct evidence that limiting IL-6 expression in endothelial cells can restore impaired VE-cadherin expression due to HIF2a deficiency.

### Therapeutic Implications of the HIF2α/ARNT Axis

The identification of the HIF2α/ARNT axis as a crucial modulator of IL6 transcription presents a potential target for therapeutic strategies aimed at reducing inflammation and preserving endothelial function in ischemic heart disease. In our study, it was found that Hif2α does not directly bind to the IL6 promoter but rather indirectly through ARNT. The role of Hif2α in regulating inflammation (Imtiyaz, Williams et al. 2010, Fan, Mao et al. 2019, Jiang, Tian et al. 2020, Macias, Moore et al. 2021) appears to be strongly dependent on cell type. This partnership of the HIF2α/ARNT axis allows it to regulate the expression of various genes, including IL6, which is crucial in the inflammatory response, which helps explain why different cells might respond differently to the same signals depending on whether or not they can form this complex.

Hypoxia is a primary pathological component in ischemic cardiovascular conditions, particularly ischemic heart disease, its effects being mediated on a molecular level largely by hypoxia inducible factors (HIFs) (Bilo, Gatterer et al. 2022). Our study aligns with the current finding of therapeutic role of hypoxia in heart diseases (Lee, Ko et al. 2019, Liu, Galli et al. 2020, Bilo, Gatterer et al. 2022) suggesting that the HIF2α/ARNT axis within the microvasculature exerts a protective role during myocardial infarction, providing a significant mechanistic insight (Bilo, 2022). Controlled hypoxia, thus, emerges as a viable therapeutic avenue. (Bilo, Gatterer et al. 2022)

Our study demonstrated that overexpression of ARNT can counteract the inflammatory and permeability effects observed in HIF2α-deficient human microvascular endothelial cells. However, further in vivo studies are needed to investigate the role of ARNT overexpression after MI. Furthermore, whereas mice carrying a non-inducible EC-specific *Hif2*α knockout mutation develop normally(Skuli, Liu et al. 2009), the EC-specific deletion of ARNT leads to liver necrosis, cardiac hemorrhage, and death in utero(Yim, Shah et al. 2006), while ARNT deficiencies in adipose cells protect mice from age and diet-induced glucose intolerance(Lee, Gesta et al. 2011), so whether the Hif2α/ARNT axis may have a role in these or other pathological processes could be investigated in future studies.

Collectively, this study demonstrates the potential therapeutic value of targeting the HIF2α/ARNT pathway for MI. Further research should focus on the broader implications of the Hif2α/ARNT axis in various pathologies and the in vivo significance of ARNT overexpression after MI. Exploring these avenues could enhance therapeutic strategies for cardiovascular diseases.

## Methods

### Data Availability

All data supporting these findings are available from the corresponding author upon request. For a detailed description of all methods and materials used, please refer to the Supplemental Methods and Major Resources Table in the Supplemental Materials.

Following left anterior descending coronary artery ligation-induced myocardial infarction, mice were randomly assigned to one of four groups: (1) sham-operated control, (2) sham-operated ec*Hif2* ^−/−^, (3) control MI, and (4) ec*Hif2* ^−/−^ MI. Any mice that did not survive the operation were α not included in the analysis of the study. To ensure unbiased analysis, all echocardiographic, cellular, and molecular assessments were conducted in a blinded manner.

### Animal

The animal research in the study was conducted in accordance with the National Health and Medical Research Council of America guidelines. We generated adult endothelial specific inducible *Hif2*α deficient mice using the CreERT2/Lox system. Experiments were performed on 10- to 12-week-old female and male mice weighing 22 to 25 g. Inducible ec*Hif2* ^−/−^ mice were generated by crossing mice carrying a loxP site flanking α exon 2 of the endothelial PAS domain protein 1 with transgenic mice under the control of the VE-Cadherin promoter. VE-cadherin-CreERT2 mice were generated by The Jackson Laboratory using sperm from the VE-Cadherin-CreERT2 mouse line, which was a gift from Yoshiaki Kubota (Keio University, Tokyo, Japan) to Dr. James Liao. *Hif2*α flox/flox mice were purchased from the Jackson laboratory. Deletion of *Hif2*α is achieved via oral administration of tamoxifen(30mg/kg) for two weeks, the system was shown to be successful in knocking out ARNT (Wu, Chang et al. 2014) Two groups served as the control groups (litter mater *Hif2*α flox/flox treated with tamoxifen or Cre, *Hif2*α flox/flox without tamoxifen).

### Genotyping

According to the manufacturer’s protocol, genetic DNA was isolated from the mouse tails using a DNA isolation kit (QIAGEN DNA isolation kit). Genotyping of *Hif2*α mice was determined by PCR amplification. *Hif2α^fl/fl^* mice were determined by PCR α amplification (forward primer: 5′-AGGCAGTATGCCTGGCTAATTCCAG-3′; reverse primer: 5′-TCT TCCATCATCTGGGATCTGGGAC-3′). Primers were also used for genotyping *VE-Cadherin-CreERT2* mice (forward primer: 5′-GCG GTC TGG CAG TAA AAA CTA TC-3′; reverse primer: 5′-GTG AAA CAG CAT TGC TGT CAC TT-3′).

### Coronary Artery Ligation-induced Myocardial Infarction Model in Mice

The myocardial infarction (MI) animal model was used as previously described with some modifications (Wu, Smeele et al. 2011). In summary, mice were anesthetized using isoflurane, with induction at 3% and maintenance at 1.5 to 2%. The animals were placed in a supine position on a heating pad to maintain normothermia (about 37C). Electrocardiogram (ECG), heart rate and respiratory rates were continuously monitored. Mice were then intubated and ventilated with a tidal volume of 200 μl and a rate of 105 breaths/min using a rodent ventilator Harvard Apparatus rodent ventilator). MI was induced by permanent ligation of the LAD coronary artery using an 8–0 nylon suture about 1_mm under the tip of the left atrium. Successful ligation was verified by pallor of the anterior wall of the left ventricle and by ST segment elevation and QRS widening on the ECG. Sham surgeries were performed using an identical procedure except no sutures on the coronary artery were placed. The chest was then closed in layers. The mice were kept warm with heating pads and were given 100% oxygen via nasal cannula. Animals were given buprenorphine for post-operative pain.

### *In vivo* blood vessel permeability Assessment

Cardiac vascular permeability in vivo was assessed with Evans Blue as described (Pei, Cai et al. 2022). To summarize, 0.5 % Evans Blue was injected into the tail veins of mice 30 minutes before sacrificing them. The level of cardiac vascular permeability was assessed by quantitative measurement of the Evans Blue incorporated per milligram of tissue in the control versus experimental mouse heart. The concentration of Evans Blue was measured spectrophotometrically at 620 nm and expressed as milligrams of dye per gram of wet tissue weight.

### Echocardiographic Image Acquisition

The echocardiography was conducted using a Visual ASonics Vevo 2100 with MS400 linear array transducer machine in mice under anesthesia at baseline, 14^th^ day, and 28^th^ day after MI as previously described(Wu, Chang et al. 2014) Animals were anesthetized with 1% isoflurane and were placed in a supine position. Both parasternal long and short-axis M-mode images were recorded, and analysis were performed. At least 10 independent cardiac cycles per experiment were obtained. The echocardiographer was blinded to the mouse genotype.

### Measurement of Blood Pressure by Tail**-**Cuff Plethysmography

To evaluate the effect of *Hif2*α deletion on blood pressure, we measured the blood pressure of mice one week after completing tamoxifen treatment and compared it to the blood pressure of control mice. We utilized a noninvasive blood pressure acquisition system specifically designed for mice and performed all measurements through tail-cuff plethysmography, as previously described (Wilde, Aubdool et al. 2017)To ensure accurate readings, mice were habituated to the measurement procedure for at least one week prior to recording baseline blood pressure values.

### Histology

After one month of MI induction, the mouse hearts were harvested under 3% isoflurane anesthesia. The hearts were then immediately fixed in 10% PBS-buffered formalin for 24 hours at room temperature. Following fixation, the hearts were dehydrated in graded alcohol, cleared in xylene, and embedded in paraffin. Serial sections of 5-μm thickness were cut using a microtome and mounted on slides. The sections were then stained with hematoxylin and eosin (H&E) to visualize the tissue architecture and Masson’s trichrome stain to evaluate the extent of fibrosis. The stained sections were imaged using an Echo Revolve microscope and analyzed using ImageJ software. The evaluation of histological sections was performed in a blinded manner by an independent pathologist/investigator of the University of Chicago to avoid observer bias.

### Cell Culture

We used Human Microvascular Endothelial Cells (HMVEC), Human Umbilical Vein Endothelial Cells (HUVEC) and immortalized human aortic endothelial cells (TeloHAEC). HMVEC cells were maintained in an endothelial cell medium from Cell biologics. HUVEC and TaloHAEC cells were maintained in M-199 medium (Invitrogen) supplemented with 20% fetal calf serum (Gibco) and buffered with 25 mM HEPES, fresh L-glutamine (final concentration, 2 mM), 100 U/ml K-penicillin G, and 100 mcg/ml streptomycin sulfate (Bio Whittaker). Culture medium was also supplemented with 100 μg/mL heparin (Sigma) and 100 μg/mL Endothelial Cell Growth Supplement (ECGS, Biomedical Technologies, Inc.). Cells were cultured in T75 flasks at 37^0^C either in 21% oxygen and 5% CO2 (normoxia) or exposed to 1% O2 (hypoxia) balanced with 5% CO2 and 95% N2 in an InVivo2400 hypoxia workstation (Ruskinn Technologies). No cell lines used in these experiments were found in the database of commonly misidentified cell lines that is maintained by ICLAC and NCBI biosample. Cell lines were routinely tested for mycoplasma contamination. Cell lines were verified to be endothelial through the presence of CD31 as quantified by qRT-PCR.

### Isolation of Murine Cardiac microvascular endothelial cells

Mouse hearts from ec*Hif2*α–/– and control mice were harvested and cut into pieces, digested with collagenase, and then labeled with CD31 conjugated magnetic beads for EC positive selection, as we previously reported (Nguyen, Zheng et al. 2021)

### Isolation of Cardiomyocytes from Mouse Hearts

Cardiomyocytes (CMs) were isolated from both control and hif2a -/- hearts to serve as controls for the endothelial hif2a deletion. The procedure adhered to the established method detailed in (Zhang and Rabinovitch 2022). Briefly, mice were euthanized in compliance with institutional guidelines, and their hearts were immediately excised and immersed in an ice-cold Ca²⁺-free perfusion buffer containing (in mM): 113 NaCl, 4.7 KCl, 0.6 KH₂PO₄, 0.6 Na₂HPO₄, 1.2 MgSO₄, 10 HEPES, 12 NaHCO₃, 10 KHCO₃, 10 Glucose, and 30 Taurine, adjusted to pH 7.4. The heart’s aorta was subsequently cannulated and then mounted onto a Langendorff apparatus for perfusion at a rate of 3 ml/min. Following a 4–5-minute perfusion with the Ca²⁺-free buffer, the solution was changed to an enzyme-rich buffer containing 0.2% collagenase type II and 0.025% protease. This continued until the heart tissue exhibited a softened consistency, approximately after 8-12 minutes. Subsequently, ventricles were minced and combined with the enzyme buffer. Through gentle agitation, cardiomyocytes were liberated, and the resultant mixture was passed through a 100 μm mesh filter. After a brief centrifugation at 20g for 2 minutes, the cells were resuspended in a stop buffer enriched with 10% FBS and incremental additions of CaCl₂ (ranging from 50 μM to a final concentration of 1 mM). The final cell preparation was suspended in M199 medium fortified with 10% FBS and 1% penicillin/streptomycin. The viability of the isolated cardiomyocytes was determined using a trypan blue exclusion test.

### siRNA Transfection

A predesigned small interfering RNA (siRNA) targeting *Hif2*α (si*Hif2*α) (Silencer Select Pre-Designed siRNA, Cat#:AM16708) and a non-targeted control siRNA (siCont) (Silencer Select Negative Control, #4390843) were purchased from ThermoFisher Scientific (Waltham, MA). to transfection, siRNAs were premixed with Dharmafect transfection reagent (ThermoFisher Scientific) in serum-free medium and incubated for 20 mins. Cells were then transfected by incubation with this siRNA mixture for 24-48 h and treated with or without 1%hypoxia for 24 hours.

### ARNT Lentivirus Construction and overexpression

We used human ARNT ORF cDNA clone (aryl hydrocarbon receptor nuclear translocator (ARNT), transcript variant 1, mRNA) in pReceiver-Lv105 vector (Genecopoeia, EX-C0312-Lv105) for lentiviral overexpression. Mutated ARNT constructs were generated in the pReceiver-LV105 vector using the Quickchange kit (Agilent) with primers designed to introduce a single S77D mutation. Lentiviral vectors for expression of ARNT, mutant ARNT, or a scramble control were packaged and titrated by the Northwestern Genomic Editing and Screening Core (GET iN Core), as previously described.(Martin-de-Saavedra, Dos Santos et al. 2022)

### Western blot analysis

Cell pellets were collected and resuspended in RIPA buffer (50 mM Tris-HCl pH 7.4, 150 mM NaCl, 1% NP-40, 0.5% sodium deoxycholate, 0.1% sodium dodecyl sulfate (SDS)) containing protease inhibitor (cOmplete Protease Inhibitor - EDTA Cocktail, Roche). 30μg of total protein lysate of each sample was separated by electrophoresis in 12% Mini-PROTEAN® TGX™ Precast Protein Gels (Bio-Rad) and transferred using a Trans-Blot Turbo System (Bio-Rad). Membranes were blocked for 1 h at room temperature using blocking buffer (5% nonfat milk in TBST) and incubated with primary antibodies at 4C0 and kept on a shaker overnight. The primary antibodies used in western blotting were IL6 (# P620, Invitrogen, 1:2000), Ve-Cadherin (# sc-9989, Santa Cruz, 1: 500), ARNT (# 5537, Cell Signaling 1:500), GAPDH (# 5174, Cell Signaling, 1: 5000), HIF1A (# NB100-479, Novus 1:500), HIF2A (# NB100-122, Novus, 1:500) while HRP-conjugated anti-rabbit IgG (# 31460, Invitrogen, 1: 10000) or anti-mouse IgG (# 62-6520, Invitrogen, 1:10000) were used as the secondary antibody. Chemiluminescence signal was developed with Clarity Max ECL Western Blotting Substrates (Thermo Fisher Scientific) and imaged on ChemiDoc XRS (Bio-Rad). Quantification of blots was performed with Image J.

### Quantitative real-time PCR (qRT-PCR)

Total RNA was isolated from cultured cell lines using TRIzol™ Reagent (Invitrogen) according to the manufacturer’s protocol. 500ng of total RNA was reverse transcribed into cDNA using iScript Reverse Transcription Supermix (Bio-Rad) according to manufacturer’s protocol. cDNA was diluted 5-fold and qRT-PCR was performed using iTaq Universal SYBR Green Supermix (Bio-Rad) in CFX Connect Real-time System (Bio-Rad). qPCR results were normalized to the expression of the endogenous control 18S. Primer sequences used in qRT-PCR analysis can be found in the indicated supplementary data.

### Immunofluorescence staining

Cells were cultured on top of glass slides and transfected with si*Hif2*α and lentivirus containing indicated ARNT overexpressing vector, and then treated with 1% hypoxia for 24h. Cells were washed twice with DPBS and then fixed with 4% paraformaldehyde for 15 min at room temperature. Slides were then washed three times with DPBS for 5 min and permeabilized with 0.250% Triton X-100 in DPBS for 15 min at room temperature. To prevent non-specific binding, cells were blocked using 20% goat serum plus 3% BSA for 1h at room temperature. Cells were incubated with indicated primary antibodies overnight at 4 °C. Cells were then washed three times with PBST for 5 mins each and incubated with secondary antibodies for 1h in the dark at room temperature. Cells were then washed three times with PBST for 5 mins each and images were taken using ECHO Motorized Fluorescence microscope. A minimum of five different images were obtained for each sample.

### TUNEL Staining

HMVEC were cultured on Lab-Tak II chamber slides and fixed with 4% paraformaldehyde. TUNEL immunofluorescent staining was performed using an In Situ Cell death Detection kit, Fluorescein (Roche), according to the manufacturer’s instructions. Images were taken by using ECHO Motorized Fluorescence microscope.

### ELISA

Blood samples from controls and *Hif2*α*^−/−^* mice were collected 12 hours after MI using the superficial temporal vein phlebotomy technique (ref). Inflammatory cytokines including IL6, IL-1β and TNFα in plasma were measured using ELISA kits (DY410-05, DY406-05, DY401-05, R&D Systems, MN) according to manufacturer’s instructions.

### Endothelial barrier function by ECIS

Endothelial cell barrier function was analyzed by an electrical cell impedance assay (ECIS). Briefly, arrays (8W10E+ Applied Biophysics, NY, USA) were coated with 1% gelatin (Sigma) for 30_minutes at 37_°C. Cells were cultured with complete cell medium until cell confluence reached 95-100%. Endothelial barrier disruption was induced by treatment with human alpha-thrombin (0.5 U/well). Endothelial barrier disruption was measured by using Electric Cell-substrate Impedance Sensing (ECIS) Zθ device (Applied Biophysics, Troy, NY). Resistance measurements were taken every 10 seconds and normalized to pre-thrombin treatment.

### Tube Formation Assay

Early passage HUVEC cells were cultured at a density of 1.2 × 10^5^ with complete growth medium in 24 well plates coated with 300 μl of Growth Reduced Matrigel (Corning, NY, United States). After 17 h, the formation of tube-like structures was visualized under a microscope. Images were taken, and the number of meshes, number of extremities, length of branches and segment length were measured and analyzed by the angiogenesis analyzer for Image J as we published before (Nguyen, Zheng et al. 2021) (Carpentier, Berndt et al. 2020)

### Apoptosis Assays

HUVEC cells were transfected with either control siRNA or *Hif2*α-siRNA in 6 well plates. Cells were then treated with or without 1% oxygen for 24h. Cells were collected and pelleted for apoptosis assays using the manufacturer’s instructions (Moxi GOII– Early-Stage Apoptosis Monitoring with FITC Annexin V and Propidium Iodide (PI)). Briefly, cell pellets were resuspended with Annexin-V Binding Buffer. 1*105 cells of each resuspended samples were further mixed with 3µl of FITC Annexin-V conjugate. Cells were incubated for 15 minutes at room temperature after mixing in the dark, followed by addition of 300 µl of Annexin-V binding buffer. Next, 2µl/ml of propidium iodide (PI) was added to the cells followed by 5 more minutes of incubation. Cells were measured for apoptosis in Moxi GO II (Orflo) using “Apoptosis (Annexin V – FITC&PI)” app.

### RNA sequencing

The total RNA was extracted from control and *Hif2 ^−/−^* heart tissues 28 days post-MI, α using RNeasy Fibrous Tissue Mini Kit (74704, Qiagen, MD) according to the manufacturer’s protocol. The RNA integrity was confirmed using a Cytation3 microplate reader (BioTek, VT). The mRNA profiling was conducted using Illimina NovaSeq 6000 sequencer by the University of Chicago Genomics Facility (Chicago, IL). The libraries were prepared using Illumina TruSeq Small RNA Sample Preparation Kit (RS-930-1012, Illumina, CA). Three biological replicates were sequenced for each treatment.

### Luciferase reporter assays

Luciferase reporter assays were used to measure IL6 promoter activity using the Secrete-Pair™ Dual Luminescence Assay Kit (GeneCopoeia) following the manufacturer’s instructions. Briefly, human IL6 dual-reporter promoter clones were transfected into cells, which expressed both Gaussia Luciferase (GLuc) and Secreted Alkaline Phosphatase (SEAP). After 48_h, the cell culture medium was collected, and luminescence was measured using a SpectraMax ID3 plate reader. The luciferase activities were normalized to the secreted alkaline phosphatase activity of the internal control.

### Chromatin immunoprecipitation

*Hif2*α and ARNT chromatin immunoprecipitation (ChIP) assays were performed using a SimpleChIP® Plus Sonication Chromatin IP Kit (Cell Signaling Technology #56383) according to the manufacturer’s instructions. We verified the quality and length of the digested fragments on 1.5% agarose gels. *Hif2*α and ARNT protein-DNA complexes were immunoprecipitated using antibodies against *Hif2*α (No. NB100-122, Novus Biologicals, USA), ARNT (D28F3, Cell Signaling Technology, USA) and normal anti-rabbit IgG (No. #3900, Cell Signaling Technology, USA). The immunoprecipitated protein-DNA complexes were purified by proteinase-K digestion, and purified DNA was then quantified by qRT-PCR using iTaq Universal SYBR Green Supermix (Bio-Rad) in CFX Connect Real-time System (Bio-Rad). Enrichment of the IL6 promoter was detected with the specific primers 5’-CTGGCAGAAAACAACCTGAACC-3’ (forward) and 5’-AGGCAAGTCTCCTCATTGAATCC-3’ (reverse) and normalized to the total input control.

### Statistical Analysis

The statistical analysis was performed using Prism software (version 8.3.0, GraphPad, San Diego, CA). Data is shown as the mean ± SEM. Data sets were determined to be normal by the Shapiro-Wilks test, and unpaired *t*-tests were used for comparisons between the two groups. For normally distributed data, two-tailed Student’s t test was used to compare two groups; For comparisons between multiple groups, a one-way ANOVA followed by Tukey’s multiple comparisons test, or a two-way ANOVA with Bonferroni post hoc test were performed depending on the number of experimental variables. A value of P < 0.05 was considered statistically significant.

## Supporting information

Title of the paper

Abbrevation

Major Resources

## Author Contributions

KU and LA carried out the experiments, with critical assistance from LL, QZ, KP, ZH, LP and AS. YL provided bioinformatics and statistical support. KU and LA wrote the initial draft of the manuscript. QS, QZ, WS, YF, DW, and JL offered critical input and edits to the manuscript. RW designed the study, wrote the manuscript, conducted animal surgeries, and supervised the entire project.

## Acknowledgments

We extend our gratitude to Dr. JK Liao for providing the inducible VE-cadherin-CreERT2 mice. The sperm used in this research was generously provided by Dr. Yoshiaki Kubota (Keio University, Tokyo, Japan) to Dr. JK Liao. We thank the Northwestern Genomic Editing and Screening Core (GET iN Core) for packaging and titrating the lentiviral vectors used for expression of ARNT, mutant ARNT, or a scrambled control, especially under the supervision of Dr. Pankaj Bhalla. Special thanks to The University of Chicago Human Tissue Resource Center (RRID:SCR_019199), particularly Terri Shihong Li, for their invaluable assistance with tissue staining. Figure 7F was crafted using BioRender.com.

