## Supplementary material for "A Novel ARNT-Dependent HIF-2α Signaling as a Protective Mechanism for Cardiac Microvascular Barrier Integrity and Heart Function Post-Myocardial Infarction": Abbrevation

**Nonstandard Abbreviations and Acronyms**

HIF	Hypoxia Inducible Factor

HIF1α	Hypoxia Inducible 1 alpha Factor

HIF2α	Hypoxia Inducible 2 alpha Factor

PHDs	Prolyl-4 Hydroxylases

PHD2	Prolyl-4 Hydroxylase-2

ECs	Endothelial Cells
ARNT	Aryl hydrocarbon Receptor Nuclear Translocator

ECIS	Electric Cell-Substrate Impedance Sensing

Il-6	Interleukin-6

IL-1β	Interleukin-1 beta

TNFα	Tumor Necrosis Factor alpha

pVHL	Von Hippel–Lindau

MI	Myocardial Infarction

ZO-1	Zonula occludens-1

HAEC	Human Aortic Endothelial Cells

HUVEC	Human Umbilical Vein Endothelial Cells

WT	Wild Type

KO	Knockout

mCMVEC	Mouse Cardiac Microvascular Endothelial Cells

hCMVEC	Human Cardiac Microvascular Endothelial Cells

BNP	Brain Natriuretic Peptide

NPR3	natriuretic peptide receptor 3

Col13a1	Collagen type XIII alpha 1 chain

CDH5	Cadherin 5

TJP1	Tight Junction Protein 1

ANGPT2	Angiopoietin 2

PLVAP	Plasmalemma Vesicle Associated Protein

ICAM-1	Intercellular adhesion molecule 1

VCAM-1	vascular cell adhesion molecule 1

VE-CAD	Ve-Cadherin

TER	Transendothelial Electrical Resistance

EF	Ejection Fraction

FS	Fractional Shortening
