## Supplementary material for "A Novel ARNT-Dependent HIF-2α Signaling as a Protective Mechanism for Cardiac Microvascular Barrier Integrity and Heart Function Post-Myocardial Infarction": Major Resources

**Major Resources Table**

In order to allow validation and replication of experiments, all essential research materials listed in the Methods should be included in the Major Resources Table below. Authors are encouraged to use public repositories for protocols, data, code, and other materials and provide persistent identifiers and/or links to repositories when available. Authors may add or delete rows as needed.

**Animals (in vivo studies)**

| **Species** | **Vendor or Source** | **Background Strain** | **Sex** | **Persistent ID / URL** |
| --- | --- | --- | --- | --- |
| Mouse | Animal Resources Center (ARC), University of Chicago | C57BL/6 | Male and female |  |

**Genetically Modified Animals**

|  | **Species** | **Vendor or Source** | **Background Strain** | **Other Information** | **Persistent ID / URL** |
| --- | --- | --- | --- | --- | --- |
| **Parent - Male** | Mouse | Animal Resources Center(ARC), University of Chicago | C57BL/6 *Hif2α^foxl/flox^* | Mice were initially  purchased from Jackson  Research Laboratory | https://www.jax.org/strain/008407 |
| **Parent - Female** | Mouse | Animal Resources Center(ARC), University of Chicago | C57BL/6 CRE | VE-cadherin-CreERT2 mice were generated by The Jackson Laboratory using sperm from the VE-Cadherin-CreERT2 mouse line, which was a gift from Yoshiaki Kubota (Keio University, Tokyo, Japan) to Dr. James Liao at the University of Chicago |  |

**Antibodies**

| **Target antigen** | **Vendor or Source** | **Catalog #** | **Working concentration** | **Persistent ID / URL** |
| --- | --- | --- | --- | --- |
| Ve-Cadherin | Santa Cruz | sc-9989 | 1: 1000 | <https://www.scbt.com/p/ve-cadherin-antibody-f-8?gclid=Cj0KCQjwspKUBhCvARIsAB2IYut91HGfacI5qpcavgi049nnMfnjIIt_c6l4L6eyDi0M4jEnOAw8VF0aAqRsEALw_wcB> |
| HIF2A | Novus | NB100-122 | 1:500 | <https://www.novusbio.com/products/hif-2-alpha-epas1-antibody_nb100-122?gclid=CjwKCAjwsMGYBhAEEiwAGUXJaTIMQ7v8ct_bp0fG4T1X_V-2KffSl4z5fjcgwAJO6BlsYiUV_F5U0hoCUpMQAvD_BwE&gclsrc=aw.ds> |
| GAPDH | Cell Signaling | 2118 | 1:2000 | <https://www.cellsignal.com/products/primary-antibodies/gapdh-14c10-rabbit-mab/2118> |
| IL6 | Invitrogen | P620 | 1:1000 | <https://www.thermofisher.com/antibody/product/IL-6-Antibody-Polyclonal/P620> |
| ZO-1 | Santa Cruz | sc-33725 | 1:100 | <https://www.scbt.com/p/zo-1-antibody-r40-76?requestFrom=search> |
| ARNT | Cell Signaling | 5537 | 1:500 | <https://www.cellsignal.com/products/primary-antibodies/hif-1b-arnt-d28f3-xp-rabbit-mab/5537> |
| Rabbit IgG | Bio Rad | 1721019 | 1:5000 | <https://www.bio-rad.com/en-us/sku/1721019-goat-anti-rabbit-igg-hl-hrp-conjugate?ID=1721019> |
| Mouse IgG | Bio Rad | 1706516 | 1:5000 | <https://www.bio-rad.com/en-us/sku/1706516-goat-anti-mouse-igg-h-l-hrp-conjugate?ID=1706516> |
| Alexa Fluor 594 Phalloidin | Invitrogen | A12381 | 1:1000 | <https://www.thermofisher.com/order/catalog/product/A12381> |
| Alexa Fluor 594 | Invitrogen | A11008 | 1:1000 | <https://www.thermofisher.com/antibody/product/Goat-anti-Rabbit-IgG-H-L-Cross-Adsorbed-Secondary-Antibody-Polyclonal/A-11008> |
| Alexa Fluor™ 488 | Invitrogen | A-21210 |  | <https://www.thermofisher.com/antibody/product/Rabbit-anti-Rat-IgG-H-L-Cross-Adsorbed-Secondary-Antibody-Polyclonal/A-21210> |

**DNA/cDNA Clones**

| **Clone Name** | **Sequence** | **Source / Repository** | **Persistent ID / URL** |
| --- | --- | --- | --- |
| ARNT ORF | NP_001659.1 | Genecopoeia | <https://www.ncbi.nlm.nih.gov/protein/NP_001659.1> |
| IL-6 promoter | [NM_000600](https://www.ncbi.nlm.nih.gov/gene/?term=NM_000600) | Genecopoeia | [Human promoter reporter clones: IL6\|NM_000600\|HPRM30562\|interleukin 6 (genecopoeia.com)](https://www.genecopoeia.com/product/search/detail.php?prt=22&cid=&key=HPRM30562&type=promoter&choose=il6) |

| **Name** | **Vendor or Source** | **Persistent ID / URL** |
| --- | --- | --- |
| HUVEC | ATCC | <https://www.atcc.org/products/pcs-100-013?_gl=1*19p37od*_up*MQ..&gclid=CjwKCAjwsMGYBhAEEiwAGUXJaUr4fS43zOShYXTa4jIUxd2qAzZFsSrMlrT2EhGYrxeaMDZhoHGk_xoCWHUQAvD_BwE> |
| TAloHAEC | ATCC | <https://www.atcc.org/products/crl-4052> |
| HMVEC-C | Cell Biologics | <https://cellbiologics.com/index.php?route=product/product&product_id=2341> |
| HEK293 | ATCC | <https://www.atcc.org/products/crl-1573> |

**Cultured cells**

**Data & Code Availability**

| **Description** | **Source / Repository** | **Persistent ID / URL** |
| --- | --- | --- |

**Other**

| **Description** | **Source / Repository** | **Persistent ID / URL** |
| --- | --- | --- |

**ARRIVE GUIDELINES**

The ARRIVE guidelines (<https://arriveguidelines.org/>) are a checklist of recommendations to improve the reporting of research involving animals. Key elements of the study design should be included below to better enable readers to scrutinize the research adequately, evaluate its methodological rigor, and reproduce the methods or findings.

**Study Design**

| **Groups** | **Sex** | **Age** | **Number (prior to experiment)** | **Number (after termination)** | **Littermates**  **(Yes/No)** | **Other description** |
| --- | --- | --- | --- | --- | --- | --- |
| Group 1 (Control-Sham) | Male/Female | 10-12 weeks | 30 | 12 | Yes | Hif flox/flox |
| Group 2  ecHif2a-/- Sham | Male/Female | 30 | 20 | 5 | Yes | ecHif2a-/- |
| Group 3  (Control-MI) | Male/Female | 10-12 weeks | 25 | 7 | Yes | Hif flox/flox |
| Group 4  ecHif-/- MI) | Male/Female | 10-12 weeks | 25 | 1 | Yes | ecHif2a-/- |

**Sample Size:** Please explain how the sample size was decided Please provide details of any a *prior* sample size calculation, if done.

**Inclusion Criteria**

1. Age and gender: 10-12 weeks at the time to study. Half Male and Half female
2. Confirmation of MI: The animals have confirmed MI by a validated method, ST segment elevation or depression, and T wave changes, to detect abnormalities that are consistent with MI.
3. Baseline characteristics: The animals have similar baseline characteristics, such as body weight, blood pressure, and cardiac function, to minimize confounding factors.

**Exclusion Criteria**

1. Death during or immediately after surgery
2. Failure of the ligation to induce a sufficient MI.
3. Signs of severe illness or distress during the experiment

**Randomization**

Randomization: The animals have been randomized to treatment groups to minimize bias and ensure validity of the results.

**Blinding**

We conducted echocardiography measurements without prior knowledge of the animals' genotyping. Additionally, we had different researchers perform the measurements, or our researchers were blinded to the experimental group assignments by concealing the identifiers. To maintain the blinding throughout the study, we recorded the data using the coded identifiers without revealing the experimental group assignments until the blinding was broken. Once the study was complete, we broke the blinding to reveal the treatment group assignments for data analysis**.**
